# Screening of potential biomarkers using activity-based protein profiling in a rat model of chemotherapy-induced peripheral neuropathy

**DOI:** 10.1101/2024.10.30.620562

**Authors:** He Miao, Yuxuan Ren, Yile Zhou, Mengmeng Miao, Tong Chen, Feng Ni, Xiaowei Chen

## Abstract

Chemotherapy-induced peripheral neuropathy (CIPN) is a debilitating condition affecting cancer patients, often resulting from the use of neurotoxic agents, such as paclitaxel. This study aimed to establish a rat model of CIPN through systemic administration of paclitaxel and to identify potential biomarkers using activity-based protein profiling (ABPP) with a desthiobiotin iodoacetamide (DBIA) probe. Male Sprague-Dawley rats received paclitaxel injections at a dosage of 2 mg/kg on Days 0, 2, 4, and 6. Behavioral assessments revealed significant mechanical and thermal pain hypersensitivity in paclitaxel-treated rats compared to control groups. ABPP analysis identified a total of 2,445 proteins and 6,765 peptides, resulting in the identification of 7 candidate proteins as potential biomarkers for CIPN. Notably, 5 proteins were significantly upregulated, while 2 protein including Q63425 (Prx) demonstrated downregulation in response to paclitaxel. These findings suggest that alterations in candidate protein levels may compromise alterations in the spinal cord in the context of CIPN. This study underscores the utility of ABPP in elucidating the proteomic changes associated with CIPN and highlights the potential of identified biomarkers like Prx for enhancing our understanding of chemotherapy-induced neuropathic pain mechanisms.

## 1. Introduction

Chemotherapy is a widely used treatment modality for various cancers; however, it often leads to a significant adverse effect known as chemotherapy-induced peripheral neuropathy (CIPN), characterized mainly by sensory alterations, although motor and autonomic impairments may also occur(Ege et al., 2024; Starobova & Vetter, 2017). Neuropathic pain symptoms, such as tingling, stabbing, and burning sensations, affect up to 68% of patients within the first month after chemotherapy, with around 30% of those treated with paclitaxel, oxaliplatin, or cisplatin continuing to experience painful symptoms six months post-treatment (Seretny et al., 2014). Paclitaxel is clinically employed to treat a range of cancers, including ovarian(Khayrani et al., 2019), breast(McGrogan et al., 2008), lung(Simon, 2014), bladder(Wu et al., 2024), esophageal cancers(Shitara et al., 2018), and anterior adenocarcinomas, as well as melanoma(Alves et al., 2018; Huang et al., 2020). Approximately 40% of patients receiving paclitaxel, a common chemotherapeutic agent, develop CIPN, significantly complicating their cancer treatment(Yan et al., 2015). This condition necessitates prolonged infusion times, dose adjustments, and, in some cases, premature discontinuation of chemotherapy, which can adversely affect treatment efficacy(Xu et al., 2022).

Understanding the pathogenesis of CIPN and developing effective treatment strategies are critical for improving outcomes for cancer patients. However, there is a notable gap in identifying and validating specific biomarkers correlated with CIPN pathogenesis, emphasizing the urgency of further research in this area.

Activity-Based Protein Profiling (ABPP) emerges as a powerful tool for addressing this gap. ABPP allows for the dynamic assessment of protein activity in vivo, enabling researchers to identify active proteins involved in disease processes(Cravatt et al., 2008). This technique offers several advantages over traditional proteomics, including the ability to detect post-translational modifications and variations in protein activity that may be key players in neuropathic pain mechanisms induced by chemotherapy(Speers & Cravatt, 2004) In this study, we utilize systemic administration of paclitaxel to establish a rat model of CIPN, which mimics the abnormal painful experience during chemotherapy. Our objective is to employ ABPP to uncover potential biomarkers that reflect underlying protein activity changes associated with CIPN. The use of a rat model ensures that findings can be translational to human conditions, providing a controlled environment to dissect the molecular pathways affected by chemotherapy. Our hypothesis posits that ABPP will delineate novel protein targets, contributing to a deeper understanding of CIPN and potentially facilitating the development of diagnostic and therapeutic interventions.

In conclusion, elucidating the biomarkers of CIPN through ABPP not only addresses the current research gap but also holds promise for enhancing patient management strategies and improving the quality of life for individuals affected by chemotherapy treatment.

## 2. Materials and methods

### 2.1. Animals

In this study, all experiments were conducted using male Sprague-Dawley (SD) rats, weighing between 200-250 g and aged 6-8 weeks, purchased from Beijing Vital River Laboratory Animal Technology Co., Ltd. The rats were housed in the animal facility at Ningbo University under specific pathogen-free (SPF) conditions, with ad libitum access to food and water. All experimental procedures adhered to the standards set forth in the Guide for the Care and Use of Laboratory Animals (NIH publication No. 80-23) as stipulated by the National Institutes of Health (NIH), and received approval from the Animal Care and Use Committee of Ningbo University.

### 2.2. Drug Treatment

Pharmaceutical-grade paclitaxel was utilized to induce neuropathic pain in the rats. Prior to administration, the rats were properly restrained, and paclitaxel was administered via intraperitoneal injection at a dose of 2 mg/kg. The injections were performed on Days 0, 2, 4, and 6 of the experiment, resulting in a total of four injections.

### 2.3. Pain Behavioral Tests

Pain behavioral tests were performed as a previous study(Pan et al., 2024). Briefly, the tests were conducted were conducted both prior to paclitaxel or saline injections (baseline) and on Day 6 following these injections. For the assessment of mechanical thresholds, the rats were individually placed in transparent plastic chambers situated on a perforated platform (Product No. 38450-278, Ugo Basile, Italy) and allowed to acclimate for approximately 60 minutes under appropriate lighting conditions. Mechanical pain thresholds were evaluated using von Frey filaments (Bioseb, USA). Each rat underwent six tests, with a minimum interval of 5 minutes between tests. The mechanical pain thresholds were calculated using the up-down method.

Thermal pain thresholds were measured using a plantar heat pain apparatus (Product No. 37370, Ugo Basile, Italy), which applied noxious thermal stimuli set at 60°C to the hind paw of the rat via infrared heat. The apparatus automatically recorded the paw withdrawal latency. A cut-off time of 30 seconds was established to prevent potential injury should the rat fail to withdraw its paw. Prior to testing, the rats were acclimated in individual transparent plastic chambers for 30 minutes. Each rat underwent three trials, and the average withdrawal latency from these trials was recorded as the final paw withdrawal latency, with a minimum interval of 5 minutes between consecutive trials.

### 2.4. Spinal Cord Sample Preparation

For rats that underwent cardiac blood collection, the right atrial appendage was excised, and a perfusion needle was inserted into the apex of the heart. The rats were perfused with cold saline (4°C) to flush the entire blood supply. Following perfusion, decapitation was performed. The rat was placed ventral side down on ice, and the fur was moistened with alcohol. The skin along the spine was incised using scissors to expose the vertebrae. The cervical vertebrae were transected, and both iliac bones were severed. Ophthalmic scissors were then utilized to open the vertebral canal along both sides of the dorsal midline, exposing the spinal cord. The L4-L6 segment of the spinal cord was carefully harvested, ensuring any attached nerve fibers were meticulously removed. The extracted spinal cord was placed into a centrifuge tube, and lysis buffer was added. Tissue homogenization was performed using an ultrasonic tissue disruptor. Following lysis, the centrifuge tube containing the sample was immediately placed on dry ice to prevent protein denaturation.

### 2.5 Synthesis and purification of the probe

The probe was synthesized and purified following a previous study(Kuljanin et al., 2021). The structure of the probe is shown in Figure 1. Reactions for synthesizing desthiobiotin iodoacetamide (DBIA) were closely monitored using proton and carbon nuclear magnetic resonance (NMR) spectroscopy along with high-resolution mass spectrometry. The final product was purified using reversed-phase high-performance liquid chromatography (HPLC) to achieve a purity exceeding 95%.

**Figure 1:**
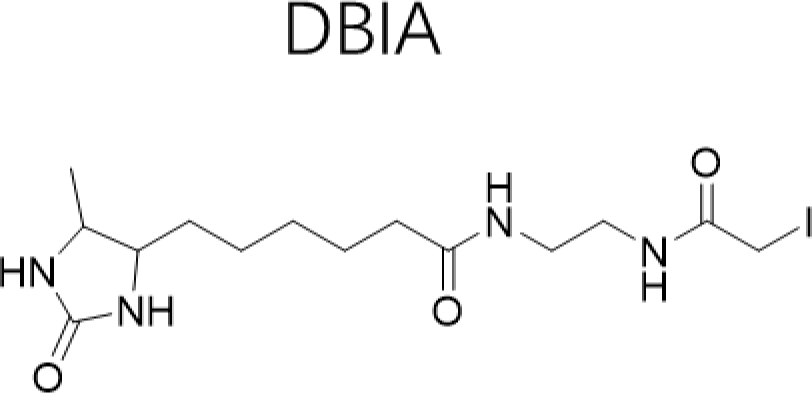
The chemical structure of the DBIA probe

To prepare DBIA, a solution containing desthiobiotin NHS-ester (100 mg, 0.32 mmol) in dichloromethane was combined with N,N-diisopropylethylamine (0.32 mmol) and N-Boc ethylenediamine (0.32 mmol). This mixture was stirred overnight at room temperature under a nitrogen atmosphere. Following the reaction, the solvent was removed under reduced pressure, resulting in a viscous oil. This oil was then treated with 20% trifluoroacetic acid (TFA) in dichloromethane (DCM) and stirred at room temperature for 3 hours. After evaporating the TFA and DCM, a yellow oil remained, which was dissolved in a mixture of 15 ml of DMF and DCM (1:2 v/v). Subsequently, N,N-diisopropylethylamine (1.6 mmol) and iodoacetic anhydride (250 mg, 0.71 mmol) were introduced. The reaction proceeded in the dark for 2 hours and was quenched by adding 10 ml of water. The aqueous layer was then separated and lyophilized before purification via reversed-phase HPLC, employing a gradient of 20–75% acetonitrile in water, with 0.1% formic acid as an additive. The fractions containing the target product were lyophilized, yielding the final compound (109 mg, 80% yield) as a white solid.

### 2.6 ABPP methods

Rat spinal tissue was lysed using ice-cold phosphate-buffered saline (PBS) and then homogenized at 4 °C. For each experiment, 200 μg of the homogenized tissue was aliquoted for downstream tandem mass tag (TMT) labeling. To assess the quantitative depth achievable with SLC-ABPP, each TMT channel containing 200 μg of cell extract was treated with 500 μM DBIA for 1 hour in the dark at room temperature. Following treatment, excess DBIA and disulfide bonds were quenched and reduced using 5 mM dithiothreitol (DTT) for 30 minutes in the dark at room temperature. The reduced cysteine residues were subsequently alkylated with 20 mM iodoacetamide for an additional 30 minutes in the dark at room temperature. To facilitate the removal of quenched DBIA and incompatible reagents, proteins were precipitated using a chloroform/methanol extraction method. Briefly, to each sample (100 μl), 400 μl of methanol was added, followed by 100 μl of chloroform, and the mixture was thoroughly vortexed. Next, 300 μl of HPLC-grade water was added to promote precipitation, and the samples were vortexed again. The samples were then centrifuged at maximum speed (14,000 r.p.m.) for 3 minutes at room temperature, and the aqueous top layer was carefully removed. The remaining pellets were washed twice with 500 μl of methanol. The protein pellets were resolubilized in 200 mM 4- (2-hydroxyethyl)-1-piperazinepropanesulfonic acid (EPPS) at pH 8.5 and digested overnight with LysC and trypsin (1:100 enzyme-to-protein ratio) at 37 °C in a ThermoMixer set to 1,200 r.p.m. The following day, the digested samples were labeled with TMT reagents or stored at −80 °C for future use.

Using the TMTpro 18plex kit (Thermo Fisher, USA), the digested peptides containing DBIA- conjugated cysteine residues were labeled with TMTpro 18-plex reagents. The reaction was quenched by incubating with 2 µl of 5% hydroxylamine for 15 minutes. All TMT-labeled samples were pooled, dried, and evaporated to dryness, and then resuspended in an enrichment buffer consisting of 100 μl of 25 mM Tris-HCl (pH 7.4), 150 mM NaCl, and 0.1% NP-40.

### 2.7 Cysteine peptide enrichment using streptavidin

Pierce streptavidin agarose beads were washed three times with phosphate-buffered saline (PBS) at pH 7.4 prior to use. To each pooled TMT-labeled sample, 100 μl of a 50% slurry of streptavidin beads was added, followed by the addition of 1 ml of PBS in a 2-ml Eppendorf tube. The samples and beads were mixed by rotating end-over-end for 4 hours at room temperature to enrich for TMT-labeled, DBIA-conjugated cysteine peptides.

To remove non-specific binding, the beads were washed using the following procedure: three washes with 0.6 ml of PBS (pH 7.4), three washes with 0.6 ml of PBS containing 0.1% SDS (pH 7.4), and finally, three washes with 0.6 ml of HPLC-grade water. To elute the cysteine-containing peptides, 500 μl of 50% acetonitrile with 0.1% trifluoroacetic acid (TFA) was added, and the beads were mixed at 1,000 r.p.m. for 10 minutes at room temperature. The eluted peptides were transferred to a new tube, and the beads were subsequently washed with an additional 200 μl of 50% acetonitrile containing 0.1% TFA, with the washes combined. The cysteine-containing peptides were dried to completion using a SpeedVac and stored at −80 °C until MS analysis.

### 2.8 Sample preparation for whole-proteome analysis

Samples were lysed with 8 M urea and 200 mM 4-(2-hydroxyethyl)-1- piperazinepropanesulfonic acid (EPPS) at pH 8.5, supplemented with protease inhibitors. The lysates were then homogenized, and DNA was sheared using a probe sonicator with a sequence of 20 pulses of 0.5 seconds each at power level 3. Total protein concentration was quantified using a bicinchoninic acid (BCA) assay, with cell lysates either processed immediately or stored at −80 °C for future use. For each TMT channel, 10 μg of protein was aliquoted for downstream processing. The protein extracts were reduced with 5 mM Tris(2-carboxyethyl)phosphine (TCEP) at room temperature for 15 minutes. Following reduction, cysteine residues were alkylated using 10 mM iodoacetamide for 30 minutes in the dark at room temperature. Samples were then digested overnight at 37 °C using LysC and trypsin (1:100 enzyme-to-protein ratio) in a ThermoMixer set to 1,200 r.p.m. The following day, the samples were labeled with TMT reagents or stored at −80 °C until further analysis.

### 2.9 Mass spectrometry

Mass spectrometry data were acquired using an Orbitrap Fusion Lumos mass spectrometer in conjunction with a Dionex Ultimate 3000 RSLCnano system. Solvent A consisted of 0.1% formic acid in water, while solvent B comprised 0.1% formic acid in 98% acetonitrile. Peptides were separated on an NTC C18 Plus column (1.7 µm, 250 mm x 75 µm, catalog number 26350-4-25S) using a 240-minute gradient: 5-25% buffer B over 200 minutes, 25-45% buffer B over 20 minutes, 45-75% buffer B over 10 minutes, 75-95% buffer B over 6 minutes, 95% buffer B for 1 minute, and 98%-2% buffer B over 3 minutes, at a flow rate of 350 nL/min. Eluted peptides were quantified using the MS3 method for TMT quantification. MS1 spectra were acquired at a resolving power of 120,000 with a maximum ion injection time of 50 ms in the Orbitrap. MS2 spectra were obtained by selecting the top ten most abundant ions for fragmentation using high energy collisional dissociation (HCD) in the ion trap, set at 35% collision energy, with a quadrupole isolation width of 0.7 m/z and a maximum ion accumulation time of 50 ms. MS3 spectra were collected at a resolution of 50,000 and a 200 ms ion injection time in the Orbitrap, with 55% collision energy, a 3 m/z isolation window for MS2, and a 0.7 m/z quadrupole isolation width.

### 2.10 Mass spectrometry data analysis

All mass spectrometry raw data were analyzed using Proteome Discoverer v.2.5 from Thermo Fisher Scientific. The spectral data were searched against a custom FASTA-formatted database that encompassed common contaminants alongside the Rattus norvegicus database (Taxon ID 10116, UniProt reviewed, 2023). The search parameters included the SEQUEST-HT search engine, a precursor mass tolerance of 50 ppm, a fragment mass tolerance of 0.9 Da, fully tryptic peptides with a maximum of two missed cleavages, and static modifications set for TMTpro18 (+304.207 Da) on lysine residues and N-termini. Dynamic modifications included DIBA probe modification on cysteine (+296.185 Da), carbamidomethylation of cysteine residues (+57.0214 Da), and oxidation of methionine residues (+15.9949 Da). Peptide spectral matches (PSMs) were filtered to achieve a false discovery rate (FDR) of 1%. For TMT quantification, only PSMs with an average signal-to-noise (S:N) ratio exceeding 3, a co-isolation threshold below 50, and an SPS mass match threshold above 65 were retained.

### 2.11 Statistics

Data are expressed as mean ± SEM. For statistical analysis, the sample size consisted of at least five animals per groups. The data distribution was determined using the Shapiro–Wilk normality test and parametric or non-parametric tests were chosen accordingly. Tukey’s post hoc tests were employed if the F-value in one-way or two-way ANOVA achieved P < 0.05. All analyses were conducted using GraphPad Prism 8 (GraphPad Software, San Diego, CA). A significance level of P < 0.05 was considered statistically significant.

## 3. Results

### 3.1 Establishment of the rat CIPN model by systemic administration of paclitaxel

We evaluated the nociceptive actions of paclitaxel in pain sensation. The schematic design is shown in Figure 2A. paclitaxel (PTX) treatment induces mechanical pain (Figure 2B, p < 0.001, PTX vs saline) and thermal pain (Figure 2C, p < 0.001, PTX vs saline) at day 6 post-paclitaxel injection.

**Figure 2.**
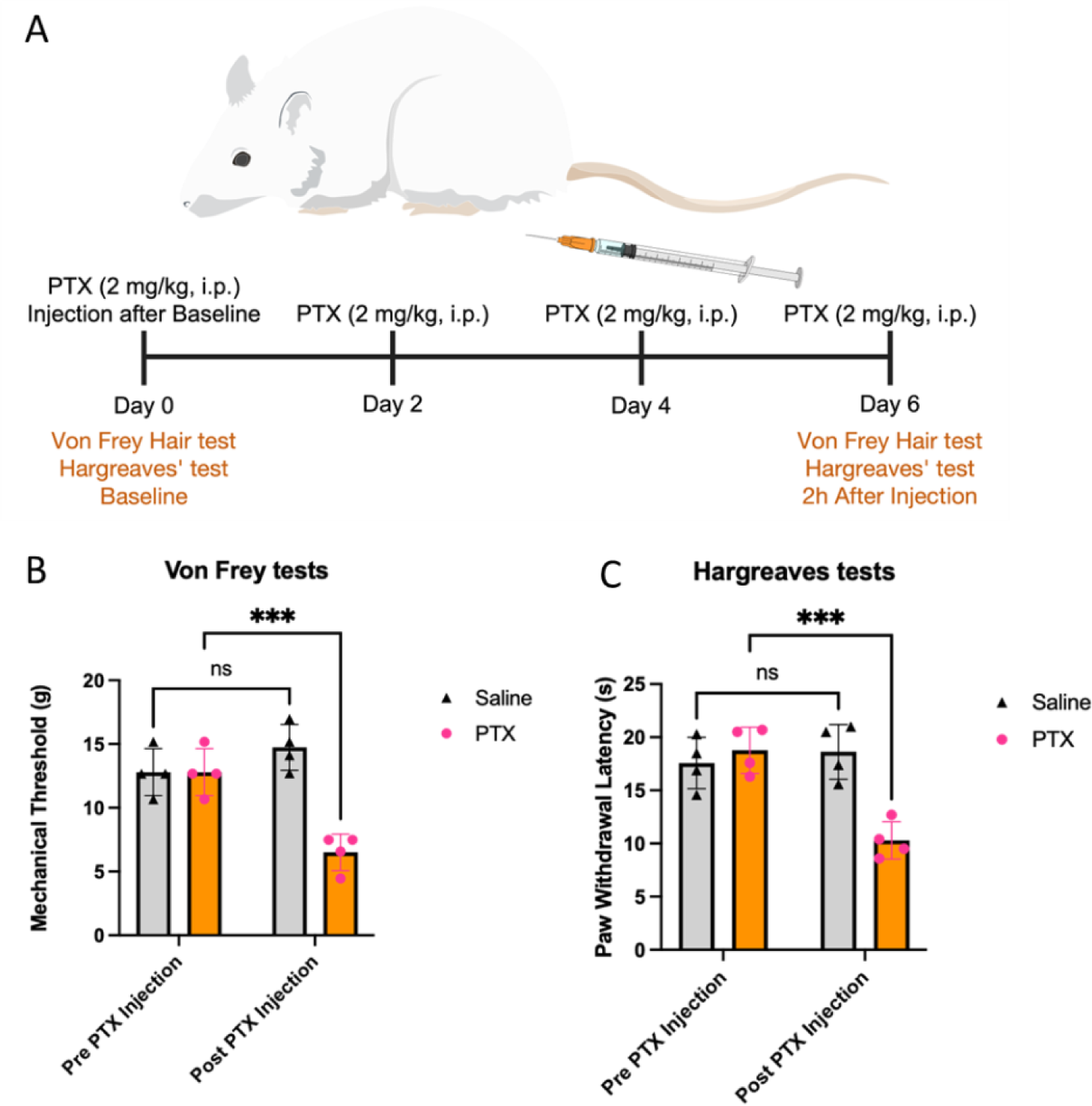
Repetitive paclitaxel (PTX) application induced mechanical and thermal pain hypersensitivity. (A) The schematic diagram illustrating the timeline of drug treatments and behavior tests. (B-C) The effect of PTX on the mechanical (B) and thermal (C) pain thresholds of paclitaxel-treated male rats. The data are expressed as mean ± SEM. Two–way ANOVA followed by Tukey’s post hoc test. ***p < 0.001 (PTX vs saline); ns, no significance.

### 3.2 ABPP Data overview and identification of candidate biomarkers

In this study, we employed the DBIA probe to analyze a total of 2,445 proteins and 6,765 peptides. Among these, 4,479 peptides exhibited quantifiable values with at least one DBIA modification, corresponding to 2,005 distinct proteins. Significant changes in protein abundance between the Saline and paclitaxel treatment groups were used as selection criteria for potential biomarkers, defined as ratios greater than 4 or less than 0.25. Through this analysis, we identified 7 candidate proteins that may serve as biomarkers for CIPN.

To assess the reliability of our data, we calculated the coefficient of variation (CV) for peptides with average quantification values exceeding 5 across the six samples in each group. The average CV was below 20%, indicating good reproducibility of the data (Figure 3).

**Figure 3:**
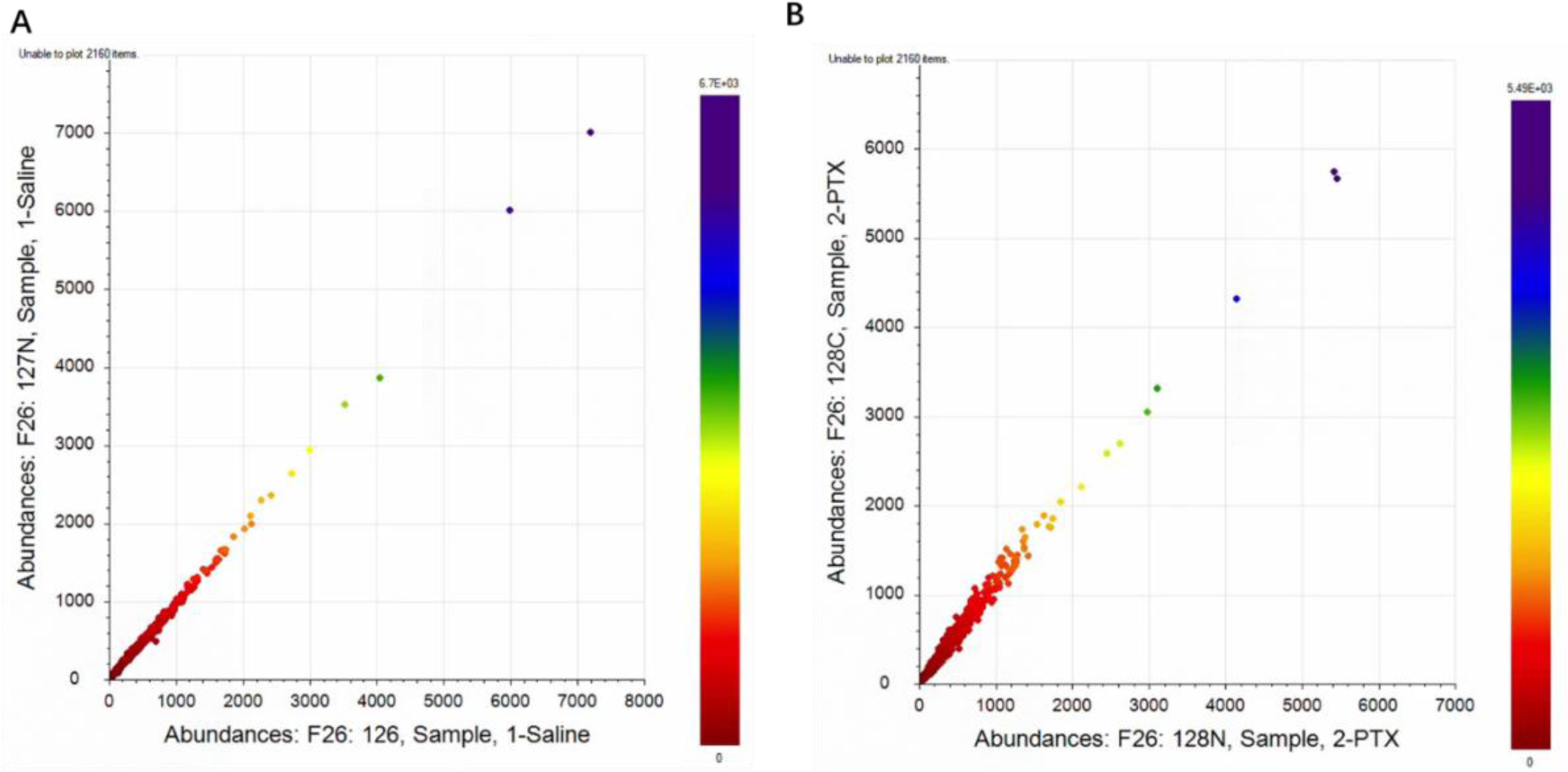
Repeatability fitting plot of two sets of samples (A for Saline; B for paclitaxel)

### 3.3 Principal Component Analysis and Cysteine Ligand Profile

We performed principal component analysis (PCA) on the six samples from each group to visualize differences in the data distribution. The PCA results, as depicted in Figure 4, revealed distinct boundaries between the Saline and paclitaxel-treated groups, confirming significant differences in protein profiles.

**Figure 4:**
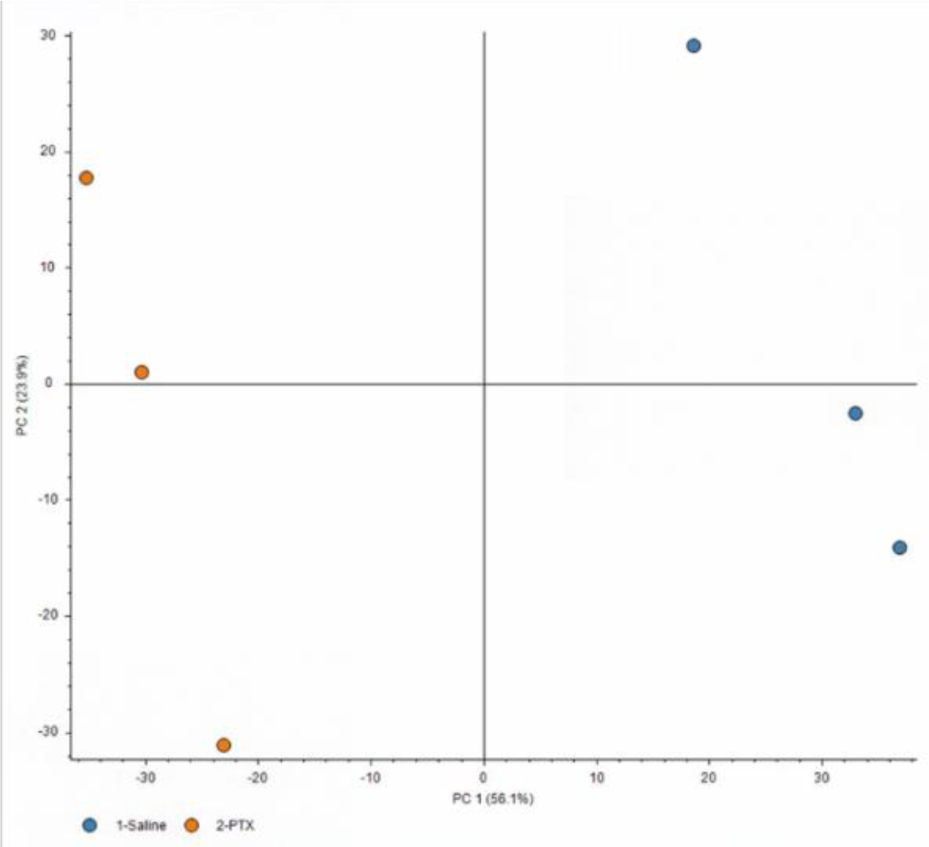
principal component analysis (PCA) on the six samples from each group

Additionally, we constructed a Cysteine Ligand Profile by plotting the abundance of DBIA-modified peptides on the x-axis against the ratio of their abundance between the Saline and paclitaxel groups on the y-axis. Volcano plots were obtained by comparing the peptide data with DBIA from the two sets of samples (Figure 5). This analysis revealed two peptide segments from proteins Q8K3P7 (5-21) and Q9JLJ3 (259-270) that were significantly upregulated in the paclitaxel treatment group, as well as two segments from proteins P02770 (311-337) and Q63425 (768-784) that were significantly downregulated compared to the Saline controls.

**Figure 5:**
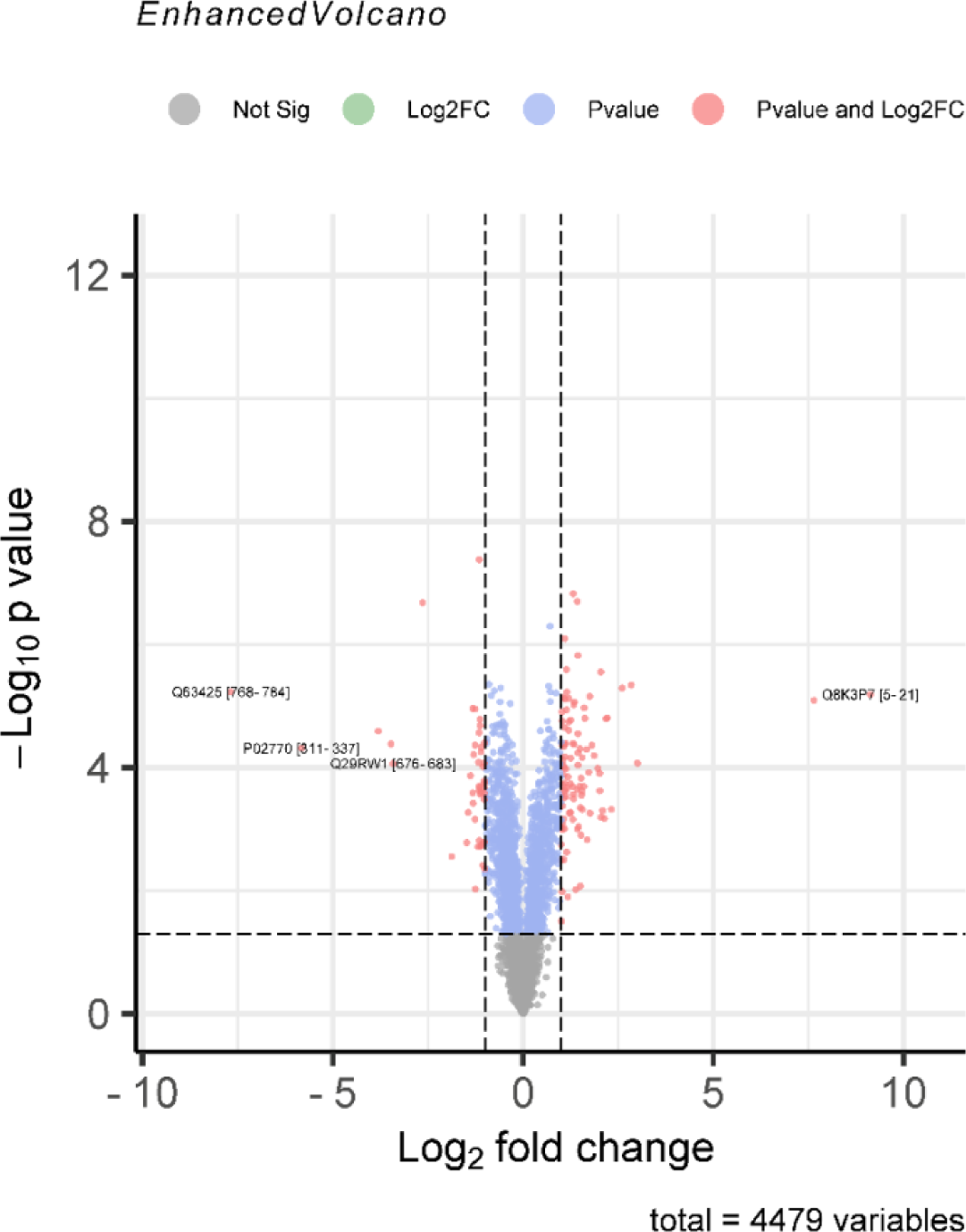
Volcano plots comparing the peptide data with DBIA from the two sets of samples.

### 3.4 Biomarker selection criteria and findings

The biomarkers were filtered based on stringent criteria: low-confidence proteins and contaminants were removed, and only peptides with at least one DBIA modification were retained. Additionally, the ratio of quantitative values between the two groups had to be greater than 4 or less than 0.25, with a corresponding p-value of <0.05. For ratios where the denominator was zero, values were adjusted accordingly for consistency.

Several proteins were excluded from biomarker consideration due to potential contamination, including P02091 (HBB1), P11517 (HBB2), and P02770 (Alb), likely related to residual blood. Non-specific binding was noted for Q29RW1 (Myh4), leading to its exclusion as well. Among the remaining proteins, Q02759 (Alox15) showed multiple peptides with consistent changes in abundance, while Q5RJP0 (Akr1b7), Q64361 (Lxn), and Q9WVK7 (Hadh) were represented by single peptides with notable changes.

Furthermore, proteins Q9JLJ3 (Aldh9a1), Q8K3P7 (Hint3), and Q63425 (Prx) were each linked to multiple peptides; however, only one peptide per protein exhibited a significant abundance change. Specifically, Q9JLJ3 and Q8K3P7 were significantly upregulated in the paclitaxel group, whereas Q63425 showed a notable downregulation. These findings suggest that paclitaxel treatment may have induced significant conformational or post-translational modifications in these proteins. Detailed information on the selected biomarkers and their characteristics can be found in Tables 1-3.

**Table 1:**
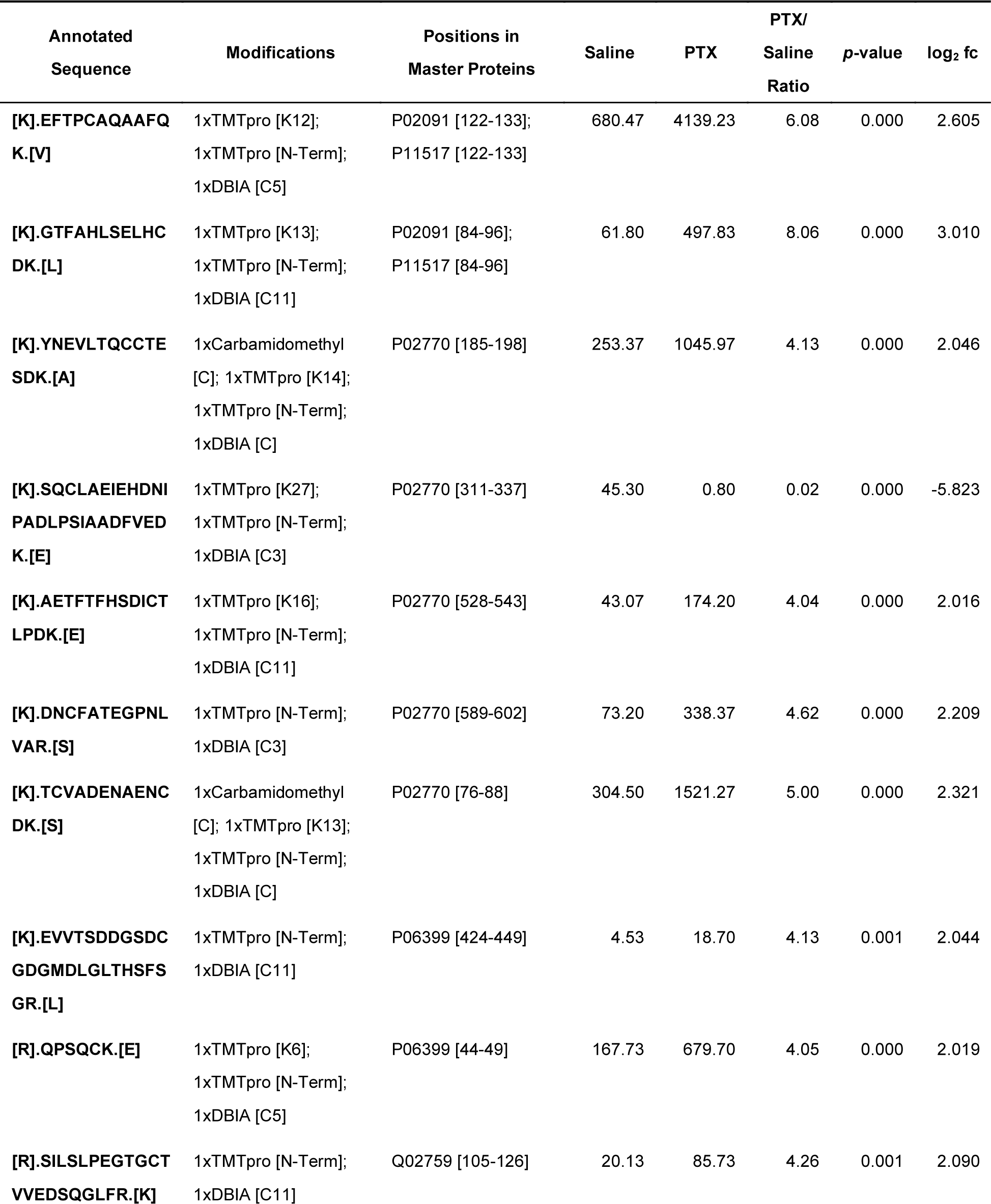
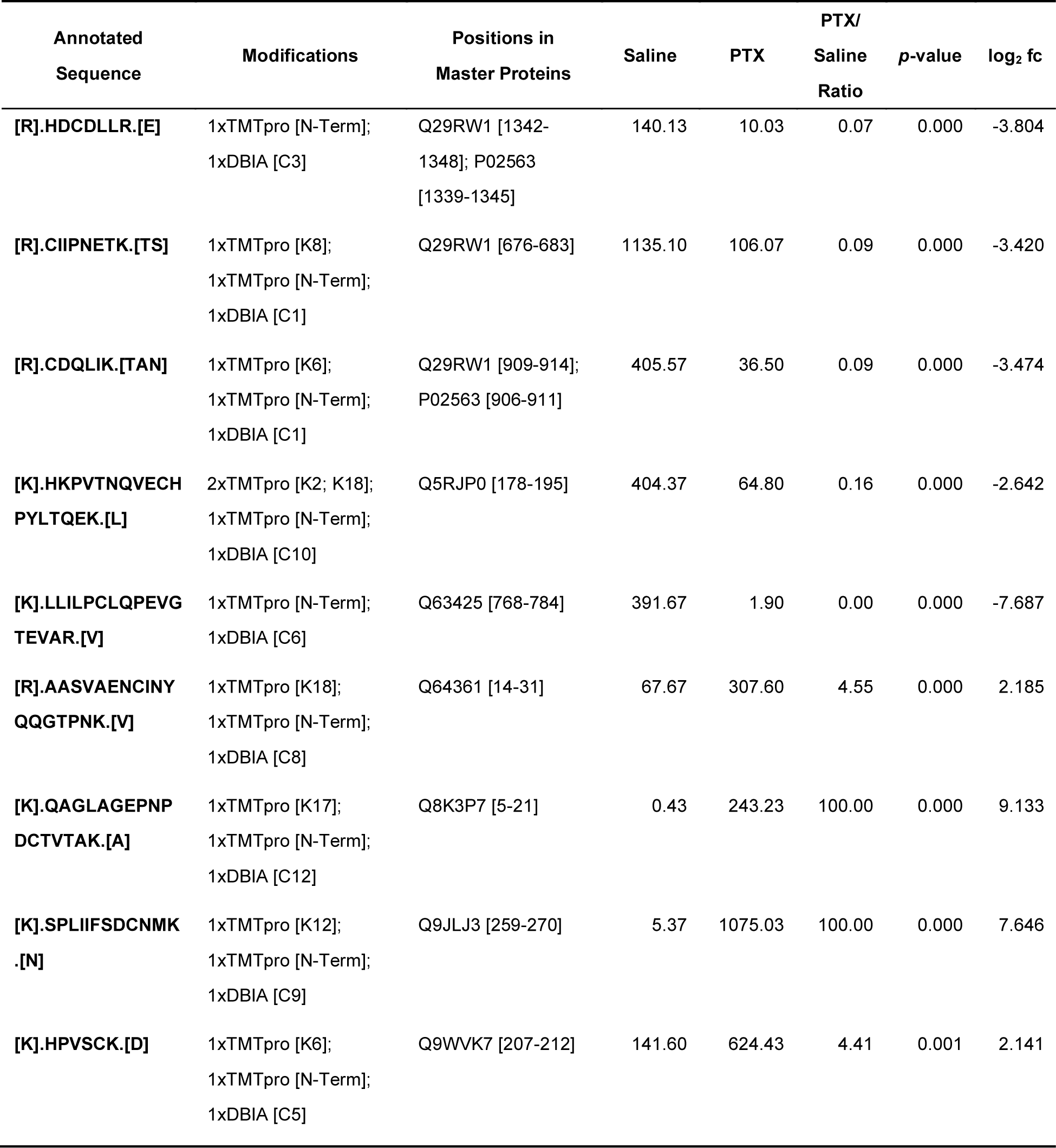
Screened target peptides by ABPP.

**Table 2:**
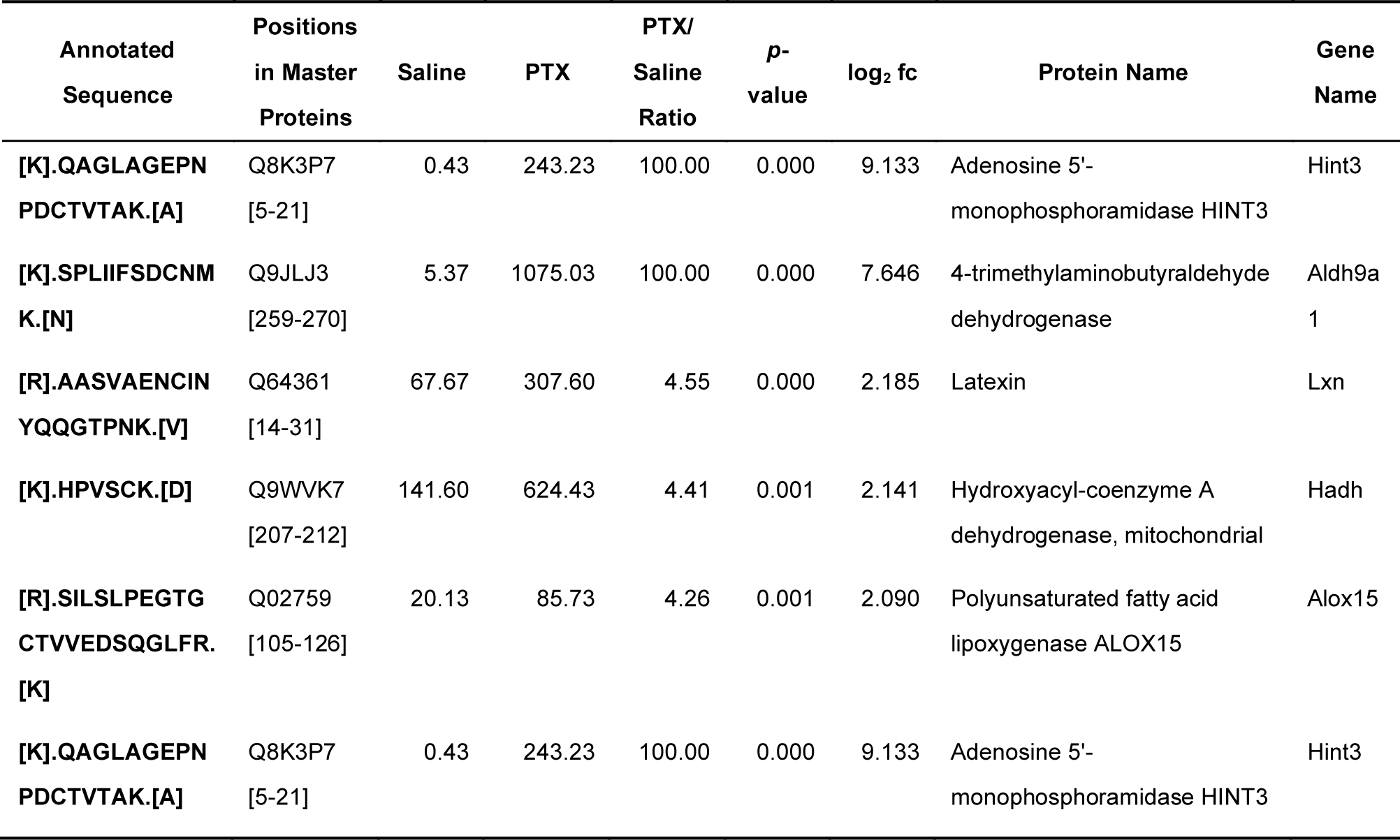
Candidate Protein Biomarkers Identified as Upregulated in CIPN.

**Table 3:**
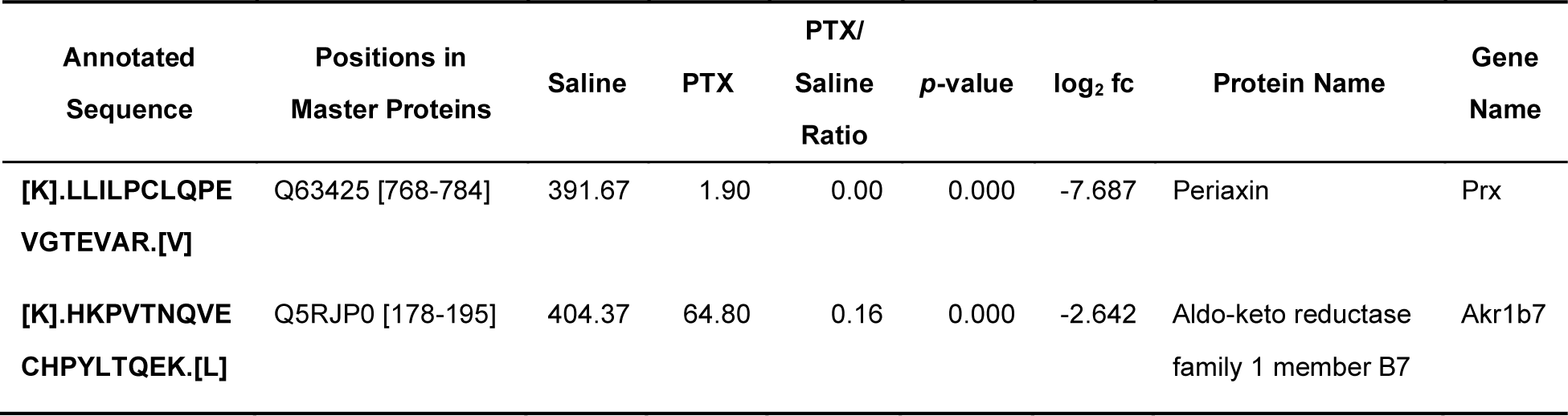
Candidate Protein Biomarkers Identified as downregulated in CIPN.

## 4. Discussion

Chemotherapy-induced peripheral neuropathy (CIPN) is a significant complication affecting cancer patients treated with neurotoxic agents, particularly paclitaxel. In this study, we established a rat model of CIPN through systemic administration of paclitaxel, leading to pronounced mechanical and thermal pain hypersensitivity. Our findings align with previous studies that documented similar effects of paclitaxel in inducing nociceptive behaviors (Mei et al., 2023; Pan et al., 2024; Xu et al., 2022). The induced pain was evaluated using behavioral tests, which demonstrated marked alterations in pain thresholds, providing a reliable model for studying CIPN mechanisms and potential therapeutic interventions.

In conjunction with the behavioral evaluations, we utilized ABPP with a DBIA probe to identify candidate biomarkers associated with CIPN in the spinal tissue. This technique enabled us to analyze a substantial number of proteins, yielding 2,445 proteins and 6,765 peptides across the samples. The ability to identify 7 candidate proteins as potential biomarkers for CIPN underscores the effectiveness of DBIA in facilitating the detection of changes in protein activity associated with chemotherapy. Among the proteins identified, specific candidates such as Q8K3P7 (Hint3) and Q9JLJ3 (Aldh9a1) exhibited significant upregulation, whereas Q63425 (Prx) showed downregulation in response to paclitaxel.

The role of periaxin (Prx) in myelination and maintenance of the peripheral nervous system (PNS) presents a compelling case for its potential as a biomarker in chemotherapy-induced peripheral neuropathy (CIPN). Prx is encoded by a gene in Schwann cells and produces two critical PDZ domain proteins essential for stabilizing myelin sheaths. The importance of Prx is underscored by a previous study that demonstrate the severe consequences of its deficiency(Gillespie et al., 2000). In Prx−/− mice, while compact PNS myelin is assembled, the sheath remains unstable, leading to demyelination and associated neuropathic pain behaviors such as mechanical allodynia and thermal hyperalgesia. The observation that these pain phenotypes highlights the underlying neural mechanisms linking Prx dysfunction, pain, and nerve damage. Notably, the loss-of-function mutations in Prx have been linked to specific neuropathies, including Dejerine-Sottas neuropathy and severe Charcot-Marie-Tooth disease(Kijima et al., 2004). The identification of non-sense or frameshift mutations in the Prx gene among patients with peripheral neuropathy. Therefore, alterations in Prx function may serve as significant contributors to neuropathy-related pathology.

Overall, our study presents novel insights into the proteomic alterations associated with CIPN induced by paclitaxel. The identified biomarkers may pave the way for further investigations into the pathophysiological mechanisms underpinning CIPN and may ultimately facilitate the development of targeted therapeutic strategies to mitigate this debilitating condition.

## Acknowledgments

We thank the technical support of the Core Facilities, Health Science Center, Ningbo University, and Ningbo University Laboratory Animal Center.

## 5. Author Contributions

Animal experiments, HM, YR, and YZ; Sample collection: HM, YR, and YZ; Analysis and interpretation of the data, and writing the manuscript: HM, YR, TC, FN and XC; Conception and design of the study, FN and XC. All authors have read and agreed to the published version of the manuscript.

